# Tensor Decomposition Discriminates Tissues Using scATAC-seq

**DOI:** 10.1101/2022.08.04.502875

**Authors:** Y-h. Taguchi, Turki Turki

## Abstract

ATAC-seq is a powerful tool for measuring the landscape structure of a chromosome. scATAC-seq is a recently updated version of ATAC-seq performed in a single cell. The problem with scATAC-seq is data sparsity and most of the genomic sites are inaccessible. Here, tensor decomposition (TD) was used to fill in missing values. In this study, TD was applied to massive scATAC-seq datasets generated by approximately 200 bp intervals, and this number can reach 13,627,618. Currently, no other methods can deal with large sparse matrices. The proposed method could not only provide UMAP embedding that coincides with tissue specificity, but also select genes associated with various biological enrichment terms and transcription factor targeting. This suggests that TD is a useful tool to process a large sparse matrix generated from scATAC-seq.

## 1. Introduction

ATAC-seq [9] is a powerful tool to profile chromatin accessibility. In particular, scATAC-seq [1] can profile the accessibility of chromatin throughout the genome within individual cells and several tools have been invented [6, 7, 29, 11, 30, 23] for scATAC-seq. Although scATAC-seq is an effective tool, data need additional information to interpret their results. scATAC-seq data can be analyzed in two ways: 1) the scATAC-seq data can be coupled to scRNA-seq data [24], or 2) traced for potential factors that bind to identified open chromatin regions from scATAC-seq data. This prevents us from understanding scATAC-seq as it is.

For example, Satpathy et al. [22] tried to find cell type-specific cis- and trans-regulatory elements, map disease-associated enhancer activity, and reconstruct trajectories of differentiation from progenitors to diverse and rare immune cell types. Giansanti et al. [8] used a predefined set of genomic regions to interpret scATAC-seq. Buenrostro et al. [4] also tried to find an association with transfactors and cis elements. Although these are only a few examples, it is obvious that they need massive amounts of external information to interpret the scATAC-seq results.

In this paper, we applied tensor decomposition (TD) [26] to the scATAC-seq dataset and found that the low-dimensional embedding obtained by UMAP applied to that obtained by TD is highly tissue-specific. This opens numerous avenues of research where scATAC-seq data can be used without additional biological information.

Although there are some studies that applied TD to scRNA-seq [20, 19], to our knowledge, there are no studies that applied TD to scATAC-seq.

## 2. Materials and Methods

Figure 1 shows the analysis flow chart. The purpose of these processes is to project individual gene expression profiles including different numbers of cells, *M_k_*, onto the common low-dimensional space, *L* ≪ *M_k_*, to concatenate multiple profiles into a matrix, to which SVD is applied to obtain a low-dimensional projection to which UMAP is applied to obtain low-dimensional embedding.

**Figure 1:**
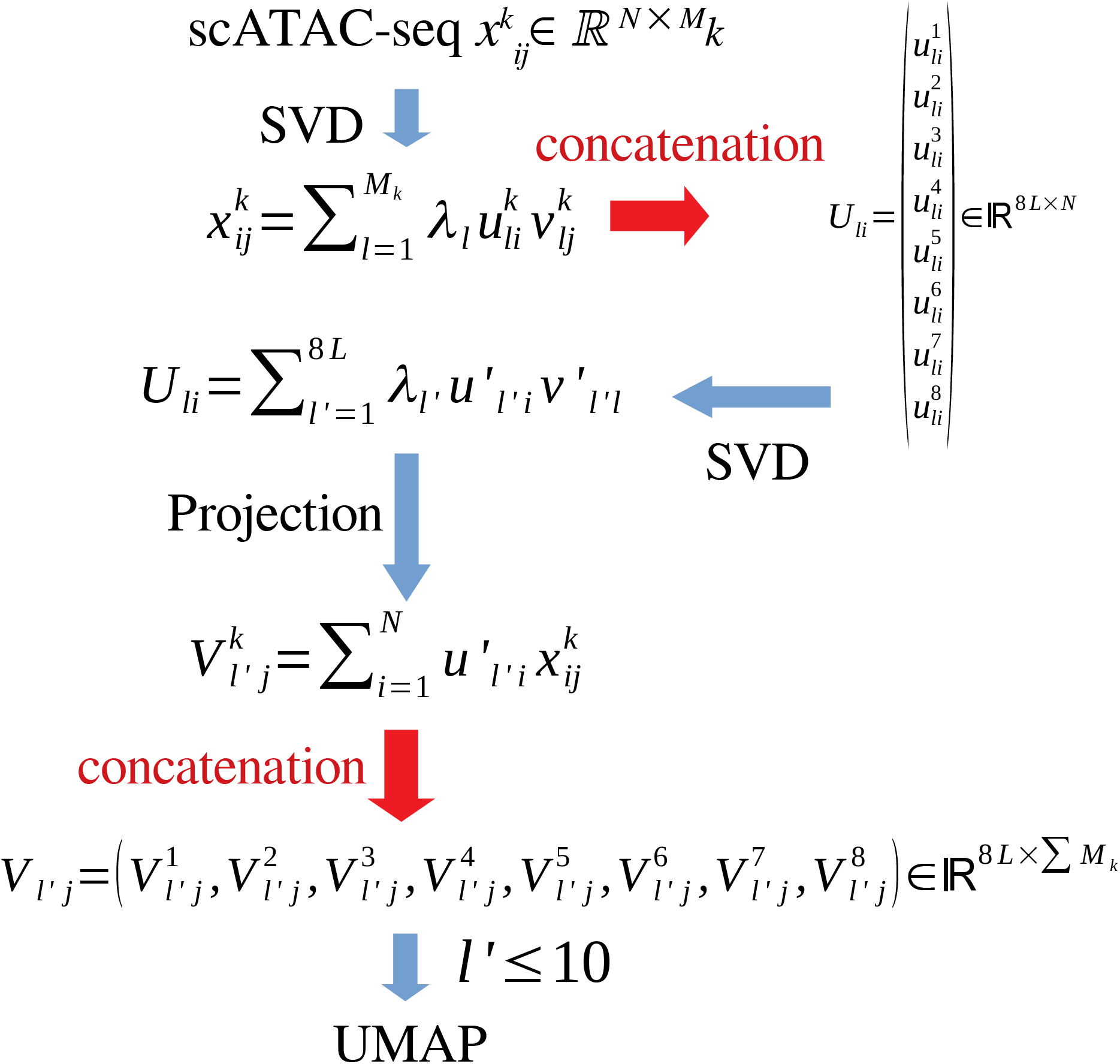
Flow chart of the analysis. 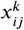 represents the expression of the *i*th gene of the *j*th cell in the *k*th set composed of *M_k_* cells. Applying SVD to 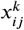, we get 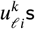, which are concatenated to get *U*_ℝ*i*_ ∈ ℝ^8*L*×*N*^. Applying SVD to *U*_ℓ′*j*_ we get 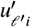, which is used to project individual 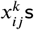 onto common space, to get 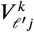, which are then concatenated to get 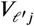, to which UMAP is applied to get low dimensional embedding.

### 2.1. scATAC-seq profiles

The scATAC-seq dataset we analyzed in this study is obtained from Gene Expression Omnibus (GEO) [3]; the ID is GSE167050 [16]. We used eight samples of GSM5091379 to GSM5091386 from four mice tissues (CTX, MGE, CGE, and LGE) with two replicates each (Table 1). Each of the eight samples is associated with three files, barcodes, features, and matrices, which correspond to single cells, genomic coordinates, and scATAC-seq values, respectively. Nuclei isolation followed the 10X Genomics ATAC nuclei isolation protocol with several modifications. All steps were performed on ice. snATAC reaction was carried out following 10X Genomics ATAC User Guide (revision C). Libraries were prepared following 10X Genomics and Illumina guidelines.

**Table 1.** Number of single cells included in the files analyzed in this study. CTX: Cortex, MGE: Medial ganglionic eminence, CGE: Caudal ganglionic eminence, LGE: Lateral ganglionic eminence

| Tissues | CTX1 | MGE1 | CGE1 | LGE1 | CTX2 | MGE2 | CGE2 | LGE2 | total |
| --- | --- | --- | --- | --- | --- | --- | --- | --- | --- |
| number of single cells ($M_k$) | 4,108 | 6,845 | 4,013 | 6,577 | 4,946 | 3,465 | 4,530 | 4,769 | 39,253 |

**Table 2.** Number of single cells included in the files analyzed in this study.

|  | Day 0 |  | Day 2 |  | Day 10 |  |  |
| --- | --- | --- | --- | --- | --- | --- | --- |
| Biological replicates | 1 | 2 | 1 | 2 | 1 | 2 | total |
| number of single cells ($M_k$) | 10,521 | 7,499 | 5,828 | 9,792 | 8,103 | 5,636 | 47,379 |

Additional dataset for validation is from ID GSE139950 [17] (Table 2). They are mice kidney of unilateral ureter obstruction at Day 0, 2, and 10. All protocols to generate scATAC-seq data on the 10X Chromium platform, including sample prep, library prep, instrument and sequencing settings.

### 2.2. Pre-processing scATAC-seq profile

The purpose of the process explained in this subsection is coarse graining output from scATAB-seq within the region whose size is comparative with that of histone. The purpose of ATAC-seq is to quantify chromosome openings, to which histone modification is believed to be critically important. Thus, it is reasonable to average the values with these regions. This procedure was employed in the previous study [27] where the integrated analysis of multi-omics single cell measurements were performed.

The values stored in the matrix files averaged over 200 bp intervals, which are supposed to correspond to the length of one wrap of chromatin and linker. Because this results in a sparse matrix, it is stored in a sparse matrix format with columns and rows equivalent to the number of single cells and the total number of intervals, which can reach 13,627,618. This value is approximately equal to 12.5 million, which is calculated by dividing the total number of mouse genome bps by the interval length (i.e. 2.5 billion by 200).

### 2.3. Application of singular value decomposition

The purpose of this section, as described in the caption for Fig. 1, is to project individual gene expression onto common space to concatenate them into a matrix to which SVD is applied to obtain a low-dimensional expression of cells to which UMAP is applied to obtain low-dimensional embedding.

To obtain a low-dimensional embedding of the individual samples *k* that will be reformatted as a tensor, we applied the singular value decomposition (SVD) to matrix, 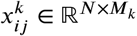 (*N* = 13,627,618 and *M_k_s* are in Table 1) as

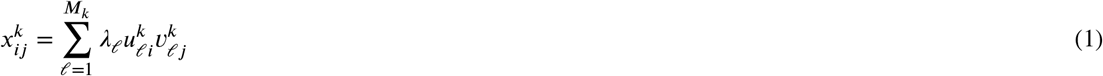

where 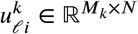 and 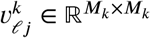 are orthogonal singular value matrices. Here *λ_ℓ_* represents the contribution of *ℓ*th singular value vectors to the individual expression profile, 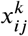. Please note that 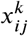 is standardized as

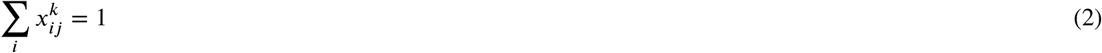

(in other words, 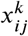 is replaced with 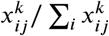) before SVD is applied. Then we get concatenated matrix *U*_ℓ*i*_ ∈ ℝ^8*L*×*N*^ of *u*_ℓ*i*_ where *L* < min(*M_k_*) is the used number of principal components. SVD is applied to *U*_ℓ*i*_ and we get

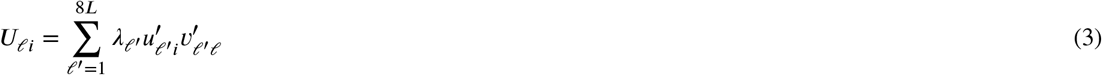

where 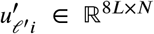 and 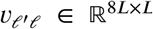 are orthogonal singular value matrices. This time, *λ_ℓ_* represents the contribution of the *ℓ*th singular value vector to all the expression profiles, not to individual ones.

Low-dimensional embedding can be obtained as

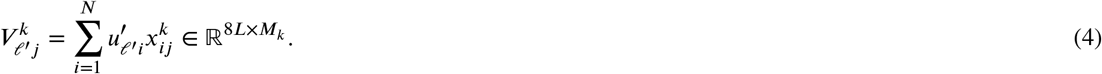

and we get concatenated matrix 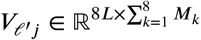

### 2.4. Sparse matrix format and singular value decomposition

Sparse matrix format is a way to store a large matrix with reduced memory by eliminating components having zero values. Because single cell measurements often have many missing values, the sparse matrix format is especially suitable for single cell measurements.

The storage of 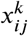 and *U*_ℓ*i*_ in a sparse matrix format allows us to deal with a matrix that has a dimension *N* greater than 10 million. It also enables us to apply SVD to these using the irlba package [2] in R [21], which is adapted for large sparse matrices.

### 2.5. UMAP

UMAP [13] is a non-linear embedding program that can embed high-dimensional data onto low-dimensional space, especially designed for preserving local cluster structures. UMAP was applied to concatenated matrix 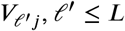 by UMAP [13] function in R with the option `umap.defaults$n_neighbors <-30` (other options remain as default).

### 2.6. Estimation of distribution of single cells in UMAP

Estimating the similarity between two single-cell embeddings is difficult because of a lack of cell-to-cell correspondence betwen two datasets. To overcome this difficulty, we proposed the computation of cell density in a local grid. If the concentration in corresponding grid points are similar, we can see that they have high similarities. To quantify the distribution of individual cells in UMAP, we divided the entire region into 10 × 10 regions. The region *S_IJ_*, 1 ≤ *I, J* ≤ 10 is {(*x, y*)|(*I* - 1)Δ*x* ≤ *x* ≤ *I*Δ*x*, (*J* - 1)Δ*y* ≤ *y* ≤ *J*Δ*y*} where Δ*x* = [max(*x_j_*)–min(*x_j_*]/10 and Δ*y* = [max(*y_j_*) – min(*y_j_*)]/10 and *x_j_* and *y_j_* are coordinates of *j*th cell in UMAP embedding. The number of single cells, *N_IJ_*, in the region *S_IJ_* is equal to the number of (*x_j_, y_j_*) ∈ *S_IJ_*. We further evaluated the correlation coefficient of *N_IJ_* between two embeddings as a measure of the similarity between two embeddings.

### 2.7. UPGMA

UPGMA is one of the hierarchical clusterings that constructs hierarchy in a bottom-up manner so that pairs associated with smaller distances are connected in a relatively lower hierarchy. UPGMA was performed with hclust function in R using option method =‘average’. The negative signed correlation coefficient of *N_IJ_* between pairs of samples was used as distance, because similar pairs (i.e., those with smaller distances) have larger correlation coefficients. Thus by adding negative signs to the correlation coefficients, they express distances.

### 2.8. Gene selection

As described in the previous study [26], we tried to select genes using 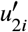. In this computation, we empirically assume that 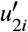 obeys Gaussian distribution. When deviation of 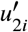 from mean value is vary large compared with standard deviation, we regard that it is too large to occur purely by chance. Although one might wonder if it is too simple, this strategy was successfilly applied to wide range of problems [26]. *P*-values are attributed to the region *i* using

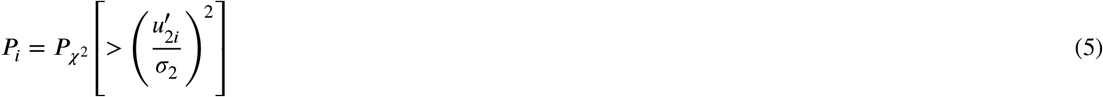

excluding *i*s with 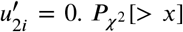 is the cumulative *χ*^2^ distribution where the argument is larger than *x* and *σ*_2_ is the standard deviation. *P*-values are corrected by BH criterion [26] and *i*s associated with adjusted *P*-values less than 0.01 are selected.

### 2.9. Genome region annotation

The selected genomic regions are evaluated by the annotate_regions function in the annotatr package [5] within Biocounductor [12]. annotatr can attribute various biological properties to individual regions.

## 3. Results

### 3.1. Obtaining low dimensional embedding

In this subsection, we aim to obtain low-dimensional embedding of cells with an integrated analysis of individual datasets. After obtaining concatenated matrix 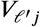 as described in the methods section, UMAP was applied to 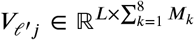 (i.e., the first *L* dimensions of 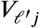 (in this case, *L* = 10)). Figure 2 shows the two-dimensional embedding of eight samples in Table 1. As observed in the investigated coordinates, the distribution of single cells in eight samples completely overlapped with one another. Nevertheless, the distribution of individual cells seems to be somewhat tissue-specific. To quantify the similarity of distributions, we divided the whole region into 10 × 10 regions and counted the number of cells in the individual regions. Figure 3 shows the scatter plots and correlation coefficients of *N_IJ_* between eight samples. The correlation coefficients between similar tissues were generally high. In few cases, correlation coefficients between different tissues were also high. To see if correlation coefficients were useful for classifying samples, we applied UPGMA (unweighted pair group method with arithmetic mean) to negatively signed correlation coefficients (Fig. 4). It is obvious that similar tissues were paired in the clustering. In addition to this, two CTX samples were clustered together, apart from six ganglionic eminence samples (LGE, CGE, and MGE), which is also biologically reasonable.

**Figure 2:**
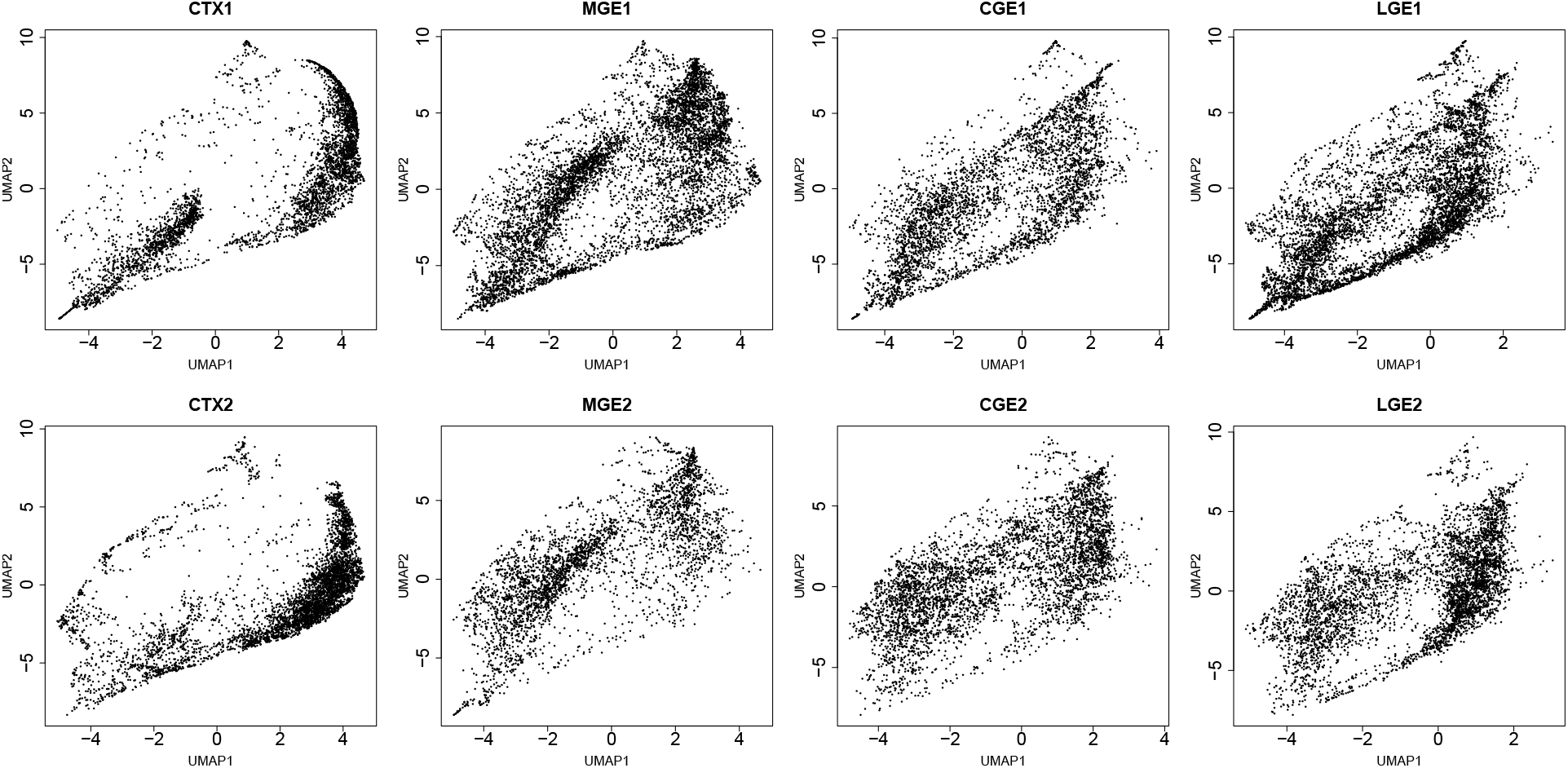
UMAP embedding of eight samples analyzed in this study (see Table 1)

**Figure 3:**
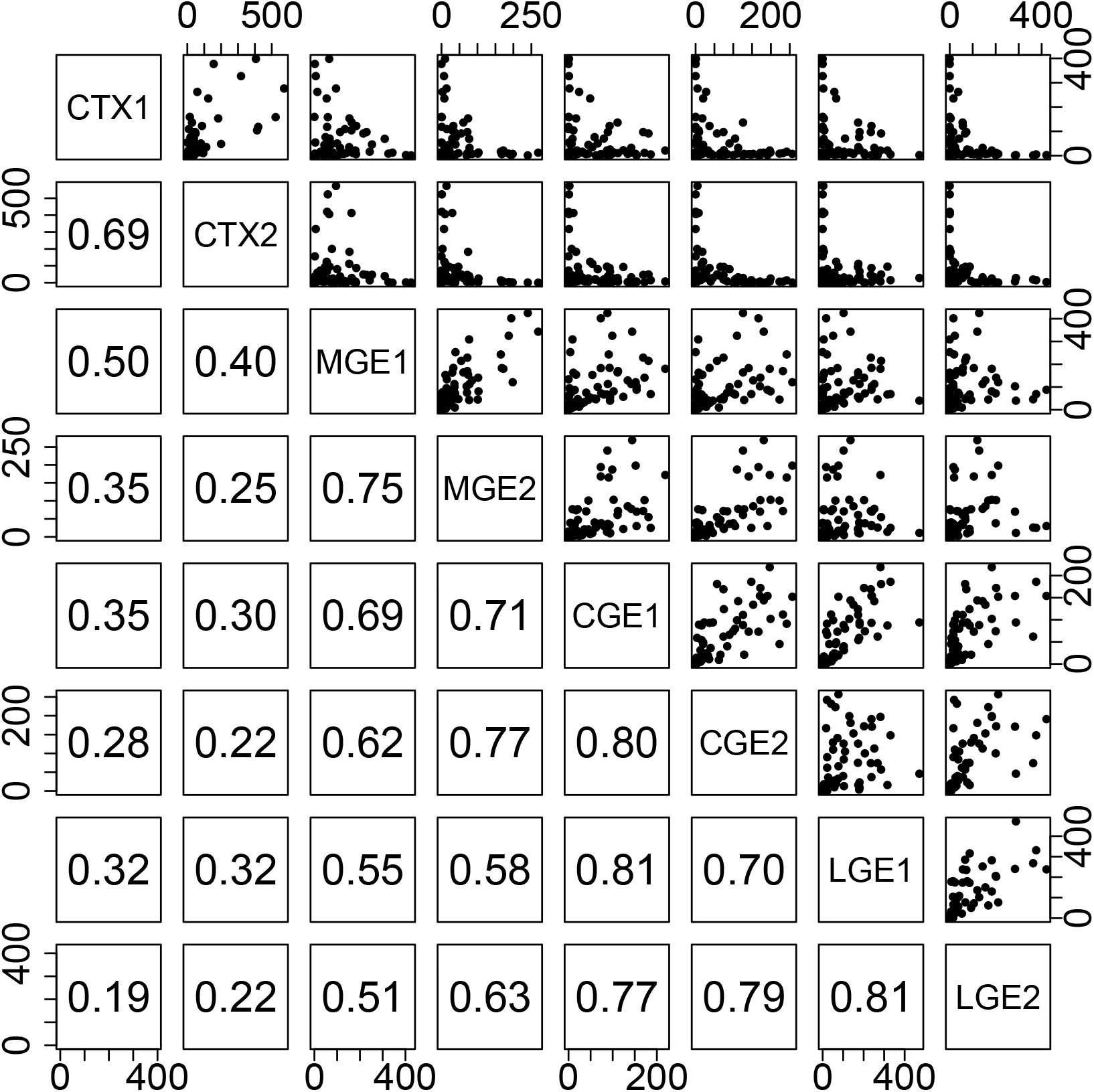
Upper: scatter plot of the number of single cells in Fig. 2, *N_IJ_*, 1 ≤ *I,J* 10, in one of 10 × 10 regions, *S_IJ_*. Lower: Kendall’s’s correlation coefficients of *N_IJ_* between eight samples.

**Figure 4:**
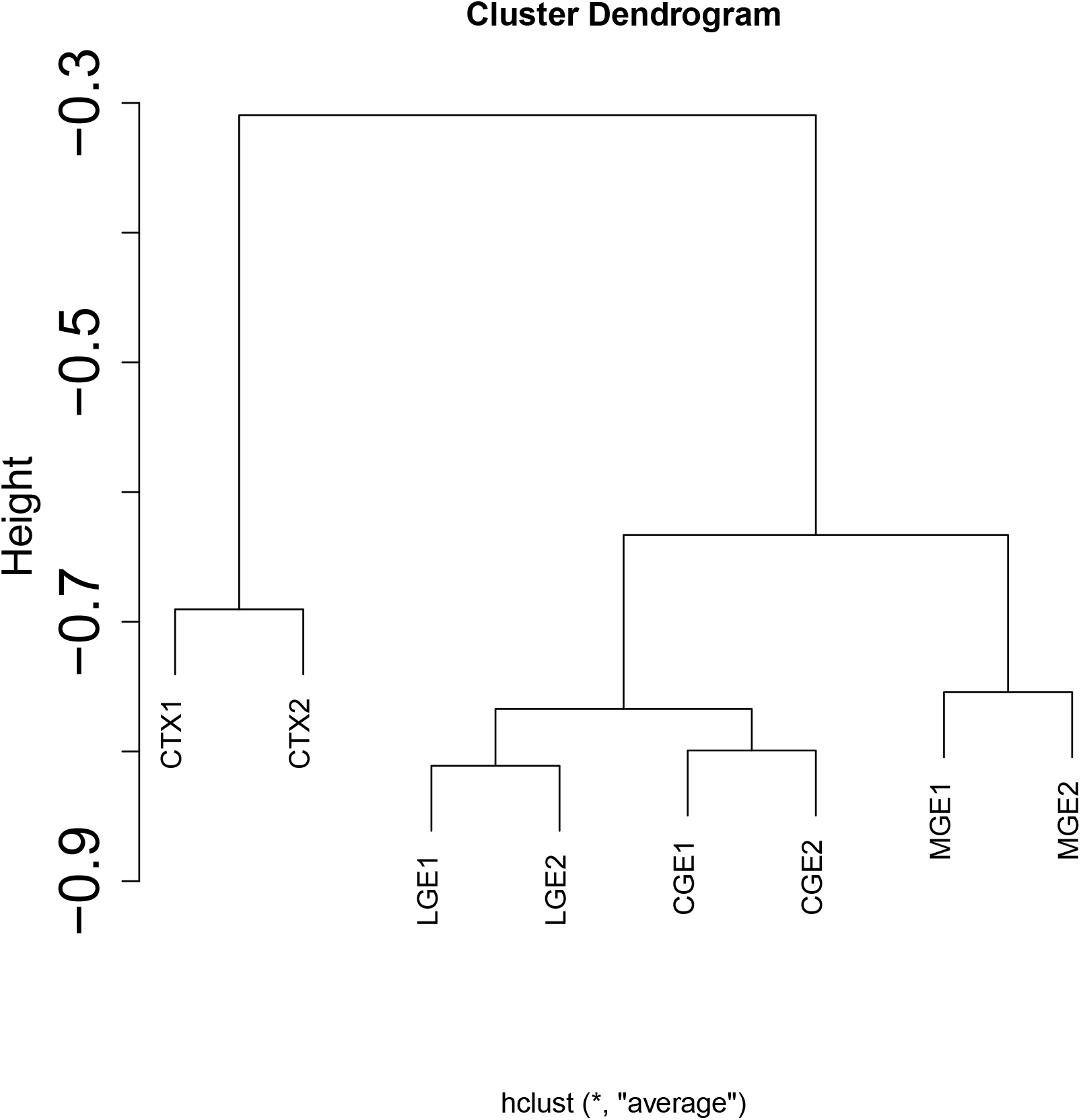
Hierarchical clustering by UPGMA of eight samples (Table 1) using coordinates obtained by UMAP. Distance is represented by negatively signed correlation coefficients in Fig. 3

### 3.2. Gene selection and enrichment analysis

In this subsection, we select genes with our own procedure and perform enrichment analyses of selected genes. Next, we tried to select genes using the results of the proposed method and biologically evaluated selected genes. We used 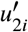 among the 10 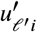 calculated to select genes using eq. (5), because the second component was often more associated with biological features in the previous studies [26]. As a result, we selected 16,469 regions. This is only 0.1% of all 13,627,618 regions.

We have biologically evaluated selected regions as in Fig. 5 and Table S1 using annotatr [5]. It is obvious that selected regions are associated with numerous functional sites in spite of the very small number of selected regions compared with the human genome (less than 0.01% as mentioned previously).

**Figure 5:**
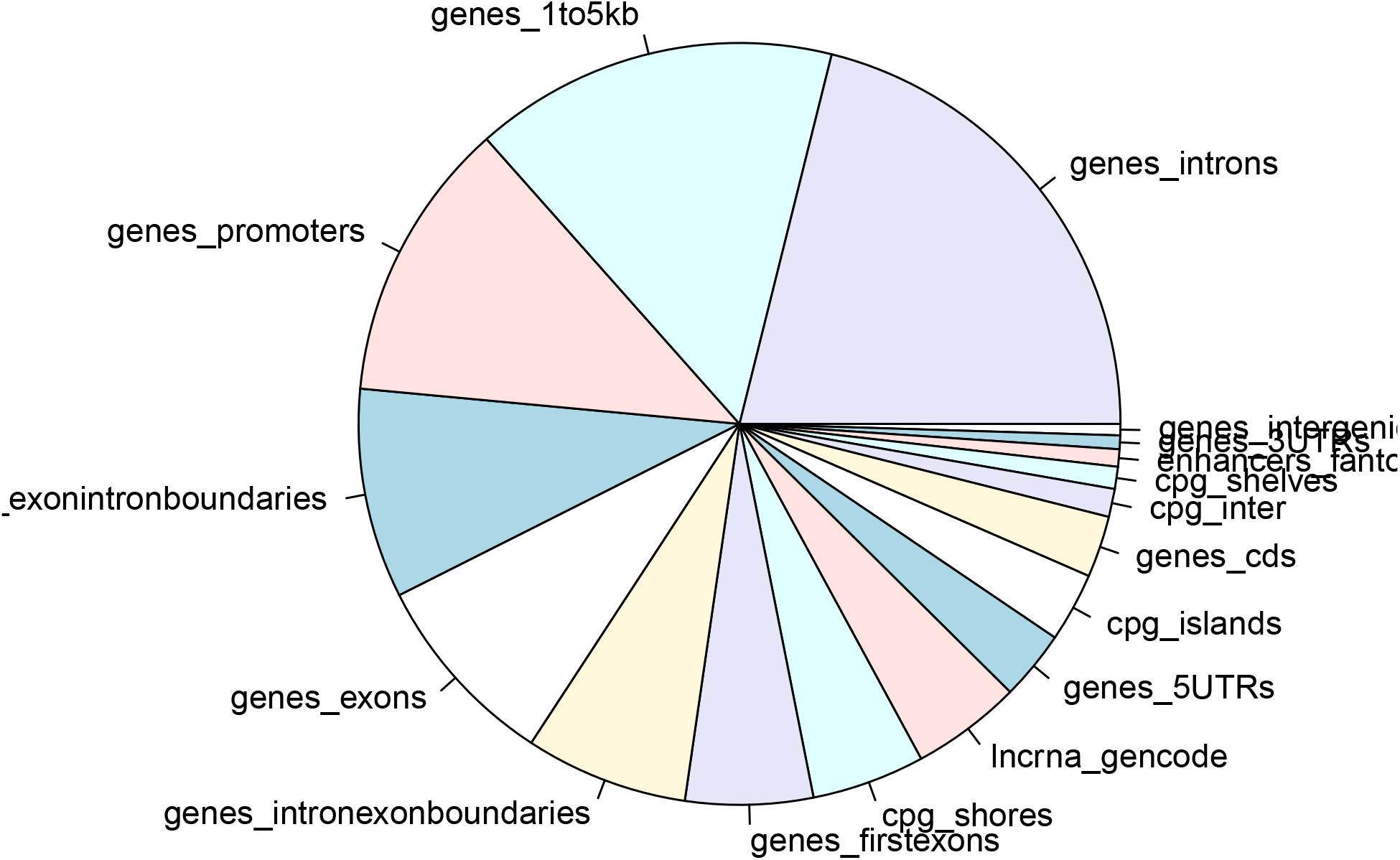
Pie chart of annotations by annotatr [5] about 16,469 regions selected by the proposed method.

To further evaluate selected regions, we uploaded 1,147 associated gene symbols identified by annotatr to Enrichr [15] (the list of 1,147 gene symbols is available as supplementary material). We found many enrichments, which are described as follows (full versions of the following tables, which list only the top 10, are available as supplementary materials). At first, transcription factors (TF) are associated with selected genes of various aspects. Table S2 lists the top 10 TFs in “ENCODE and ChEA consensus TFs from ChIP-X” in Enrichr. As can be seen, the *P*-values are very small, and thus the results are very significant. Because scATAC-seq is supposed to detect open chromatin to which TFs bind, this is reasonable. Table S3 lists more enrichment of TFs. Not only are *P*-values as significant as in Table S2, but some TFs, IRF3, SP1, and SP2, are commonly selected. Table S4 also lists additional enrichment with TFs. In contrast to Tables S2 and S3 that are based on experiments, Table S4 is sequence (motif) based. It still has highly significant enrichment of TFs, although its significance steadily decreases. The results listed in Tables S2, S3, and S4 coincide with the fact that the scATAC-seq detects open chromatin to which TFs bind.

Next we consider tissue specificity. Even if selected genes are associated with TF target genes, if it is not related with tissues where HTS was performed, it is not trustable. Table S5 lists the top 10 experiments in “Allen Brain Atlas 10x scRNA 2021” in Enrichr. It is obvious that selected genes are also associated with tissue specificity. It is interesting to note that “Allen Brain Atlas” does not consider that single cell is not coincident with the selected genes (not shown here). This suggests that we have to take into account whether it is taken from bulk cells or single cells when we consider tissue specificity.

Although Table S6 also lists the associated brain tissue specificity, some other tissue specificity is also associated. Table S7 is full of transcription activities, and Table S8 is full of DNA binding. It is also coincident with the fact that scATAC-seq detects open chromatin.

All of these analyses suggest that the selected genes are biologically reasonable.

### 3.3. More about tissue specificity

In this subsection, we focus how we can make use of cell distribution in low-dimensional space to identify the differences among individual tissues. Although we have made use of scATAC-seq to investigate tissue specificity, it is preferable to discuss distinctions in the cell diversity among individual tissues. In our analysis, distinctions among the cell diversity between individual tissues should be reflected in the distribution of single cell embedding in Fig. 2. Thus, we selected top 10 *N_IJ_*s whose difference between CTX1 or CTX2 and MGE1 or MGE2 is large. Figure 6 suggest *N_IJ_*s selected are coincident among pairs of CTX and MGE. Thus, these likely represent single cell-level difference between tissues. To understand biological aspects, we investigated which 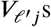 are most distinct between single cells in (*I, J*)s associated with the top 10 *N_IJ_*s. Then we found that *ℓ* = 10 and 7 correspond to the cases CTX > MGE and CTX < MGE, respectively. Then using the corresponding 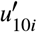 and 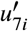, we selected 3,138 and 2,510 regions, respectively. Downstream analyses applied to the 16,469 regions in the above were repeatedly applied to these 3,138 and 2,510 regions, too. Figure 7 shows the annotated pie chart. It is obvious that it differs from Fig. 5. Accessible genomics regions enhanced in Fig. 7 are more enriched with introns and intergenic genomic regions. This suggests that cell-type specific genome accessibility is more likely associated with a non-exon or non-gene body. Enrichment analyses (see supplementary materials) also show the clear difference from 16,469 regions. For example, GO BP for CTX > MGE is enriched with synaptic functions. It is reasonable since CTX is more likely associated with synaptic activity than MGE. This means that distinct distribution of single cells in embedding between tissues can help us to understand single-cell level diversity as well. To our knowledge, no other methods could make use of the difference of single cell distribution in embedding to characterize the single-cell level diversity.

**Figure 6:**
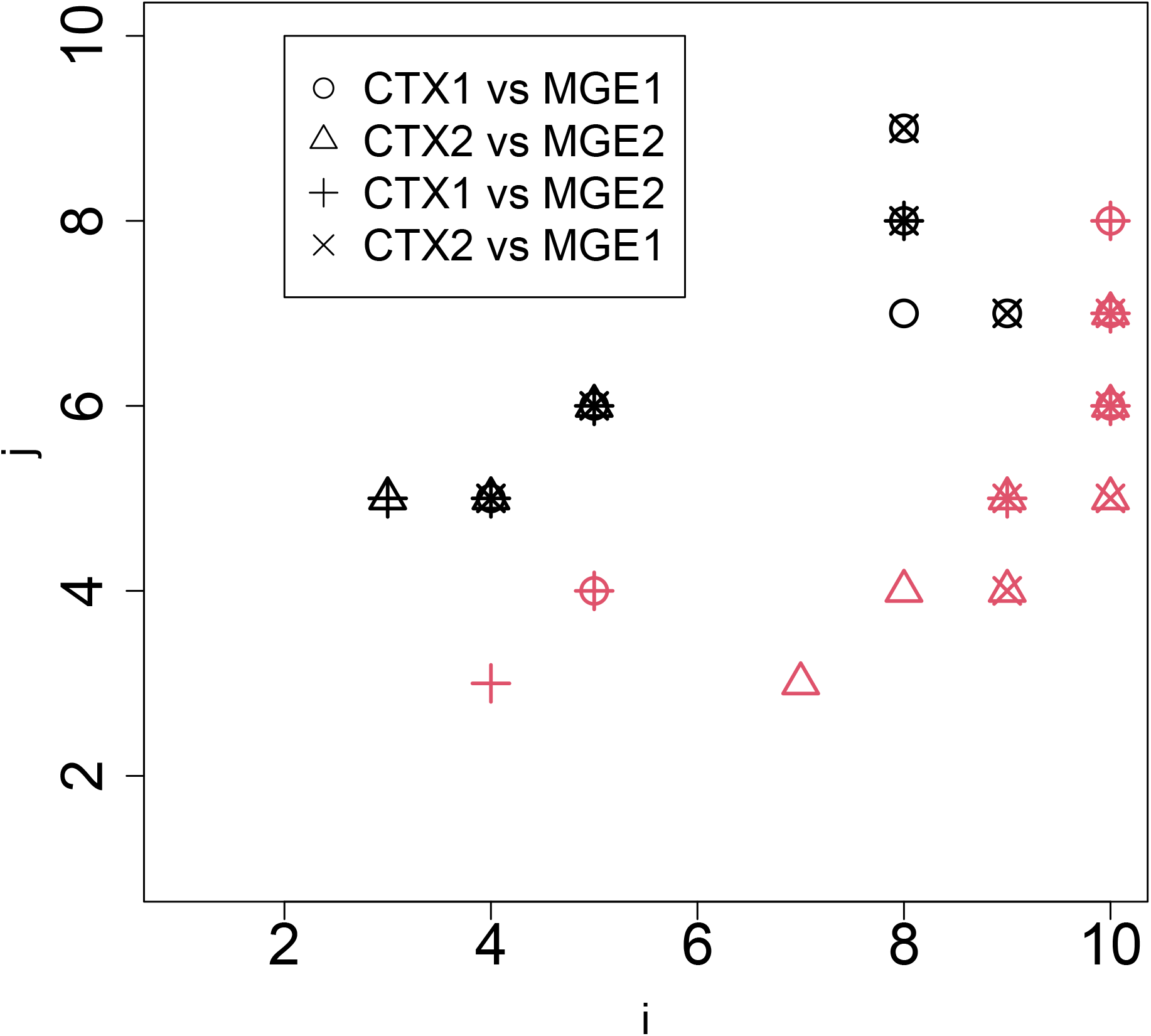
Top 10 *N_IJ_*s whose difference between CTX1 or CTX2 and MGE1 or MGE2 is large. Red and black marks correspond to CTX1 or CTX2 larger than MGE1 or MGE2 and CTX1 or CTX2 less than MGE1 or MGE2, respectively. Horizontal and vertical axes represent *I* and *J*, respectively.

**Figure 7:**
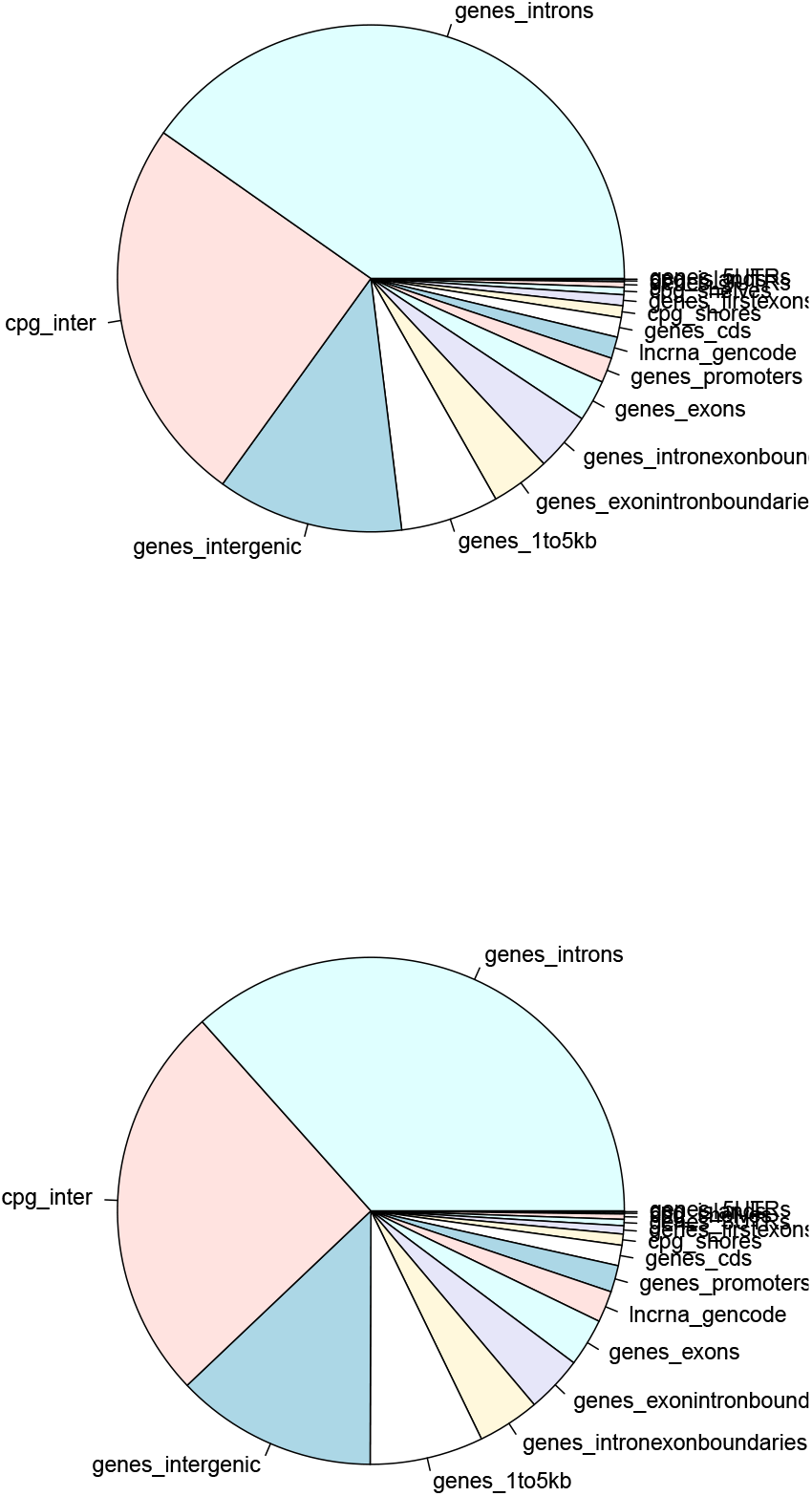
Pie chart of annotations generated as in Fig. 5, but for 3,138 and 2,510 regions associated with *N_IJ_* of CTX > MGE (upper) and CTX < MGE (lower), respectively.

## 4. Discussion

### 4.1. Absence of comparative methods

Although TD can generate the feature that can cluster samples properly (Fig. 4), if other methods cannot do this, the proposed method is more efficient and unique in terms of tensor representation. Currently, there are very few tools to process scATAC-seq data with only matrix data. For example, although the extended data in Fig. 1 of the past study [10] summarizes 10 *de facto* standard methods that can work with a scATAC-seq dataset, no methods can process the scATAC-seq dataset with only matrix data. We also tried some methods [28, 18, 14, 25] that are not included in the above list. No tools could efficiently process 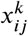 efficiently, but it is possible in spite of the large *N*. Thus, the proposed method is the only one that can work with a dataset of this size. One of the reasons the proposed method can handle this large *N* is that it stores the dataset in a sparse matrix format. SVD performs using the tool adapted for the sparse matrix format, and we do not need to process dense format. Thus, the proposed method can deal with huge datasets. Of course, this is not the only reason; one of the most popular tools, signac, can accept sparse matrix format but cannot process this of a large dataset (because it does not have not enough memory).

### 4.2. Superiority over SOTA, signac

To further confirm the inferiority of signac to our method, we applied signac to two pairs of samples, CTX1 and CTX2, as well as CTX1 and MGE1, because signac was unable to process the eight samples at once (as previously mentioned), although signac could accept a sparse matrix format in contrast to other methods specific to the scATAC sequence. Figure 8 shows the results when two signac-implemented strategies, merge and integration, were applied to the two pairs of samples, respectively. It was obvious that signac did not recognize that CTX1 and CTX2 were the same tissue, since CTX1 and CTX2 do not overlap and are completely separated. In reality, the separation between CTX1 and CTX2 is similar to that between CTX1 and MGE1 and are not the same tissues. In addition to this, in contrast to CTX1 and CTX2 in Fig. 2, where they look similar, those in Fig. 8 do not look similar. Thus, ours is better than signac in recognizing the identities of the sames tissues as well as distinguishing between different tissues using only the scATAC-seq dataset.

**Figure 8:**
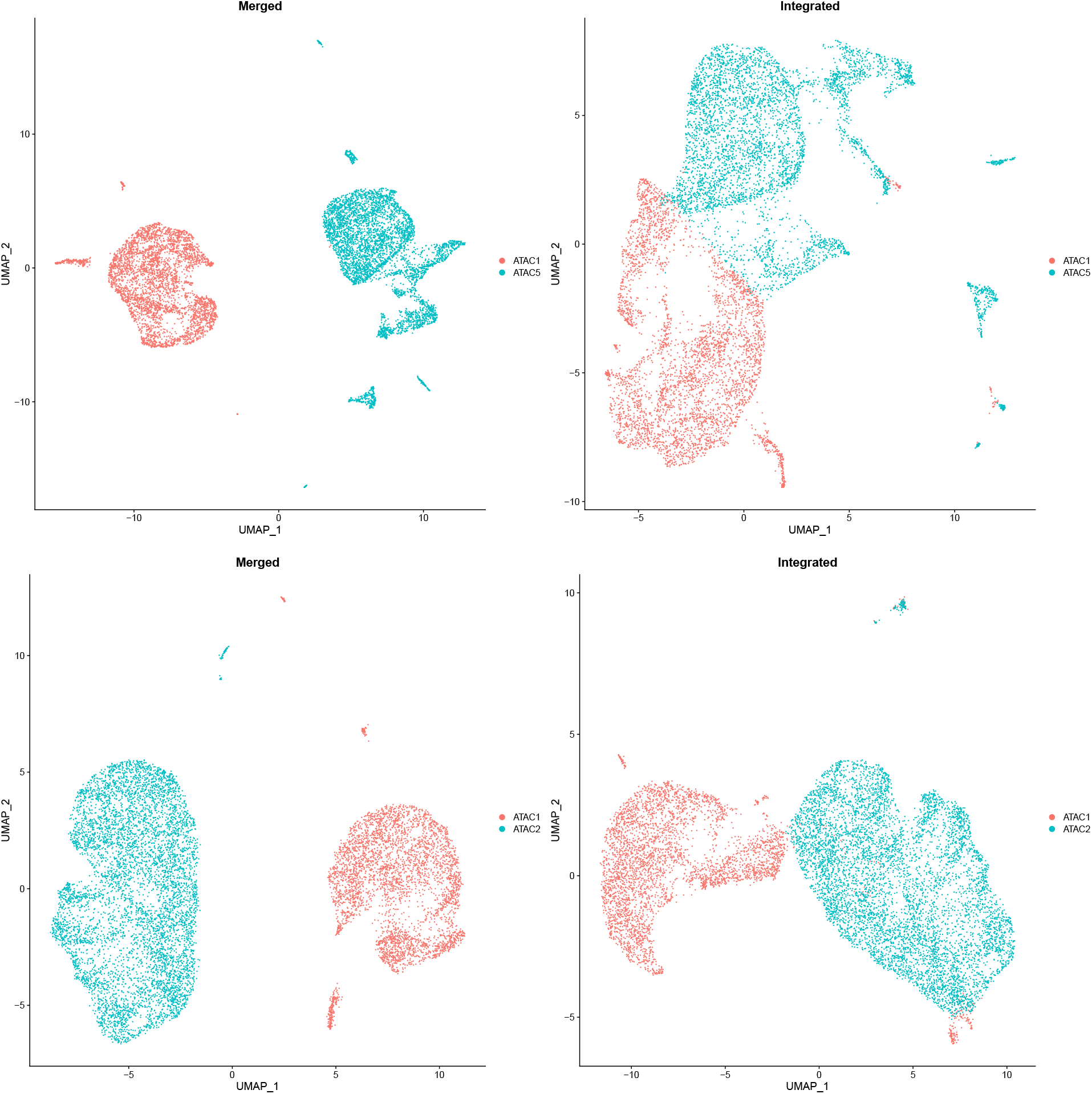
Upper: UMAP representation of signac applied to the pair of CTX1 and CTX2 (ATAC1(orange) and ATAC5(cyan) correspond to CTX1 and CTX2, respectively). Lower: UMAP representation of signac applied to the pair of CTX1 and MGE1 (ATAC1(orange) and ATAC2(cyan) correspond to CTX1 and MGE1, respectively).

### 4.3. Investigation of an additional dataset

One might also wonder if the excellent performances are accidental and not reproducible if other datasets are considered. To deny this possibility, we considered another dataset and repeatedly applied the procedure to it (Figs. 9, 10 and 11). It is obvious that embedding is coincident between biological replicates but distinct between non-biological replicate samples. Thus it is unlikely that tissue-specific embedding is accidentally obtained specific to only one set of samples. Our methods are robust and are expected to be applied to wide range of samples for scATAC-seq.

**Figure 9:**
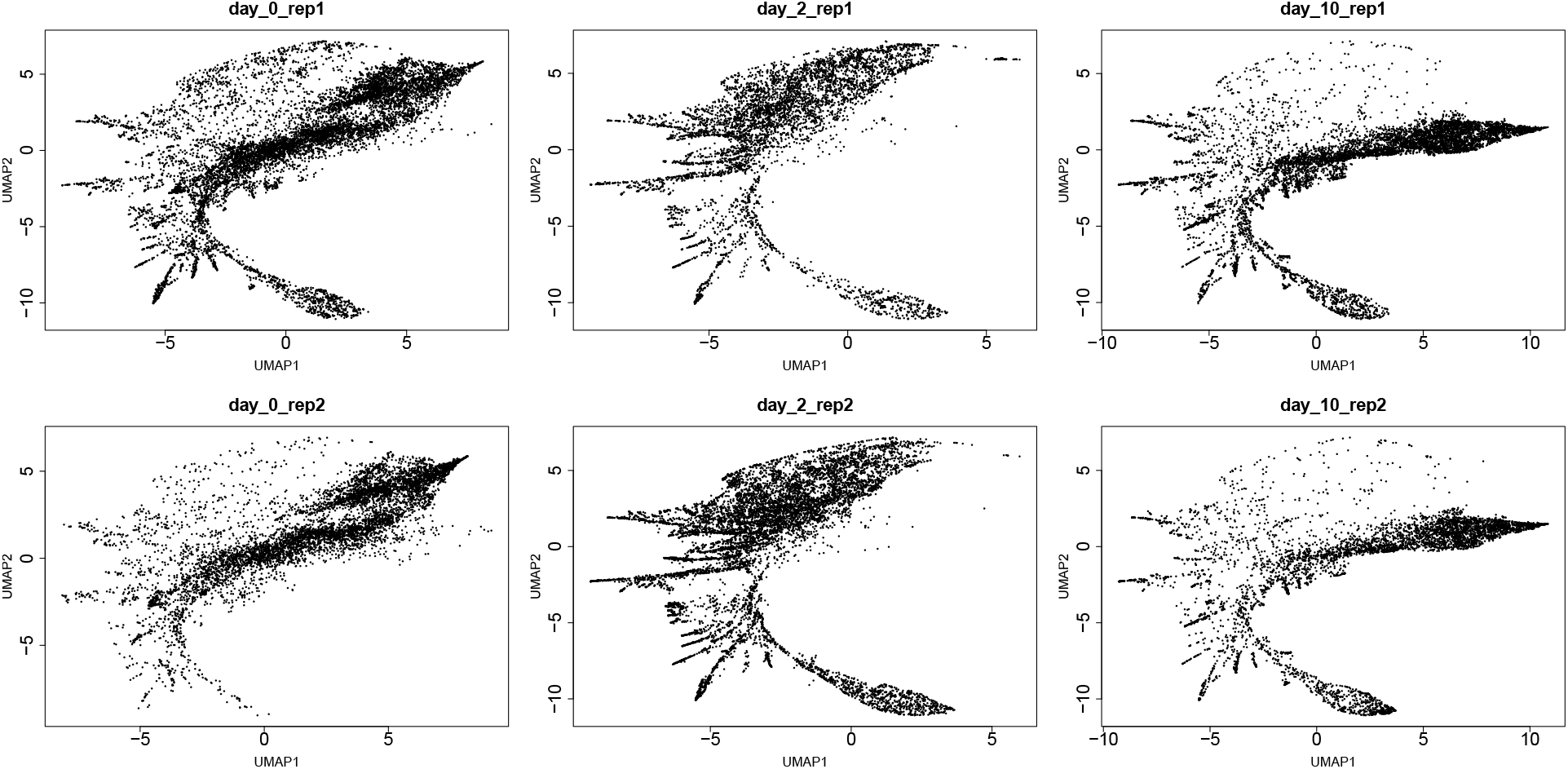
UMAP embedding of six samples analyzed in this study (see Table 2)

**Figure 10:**
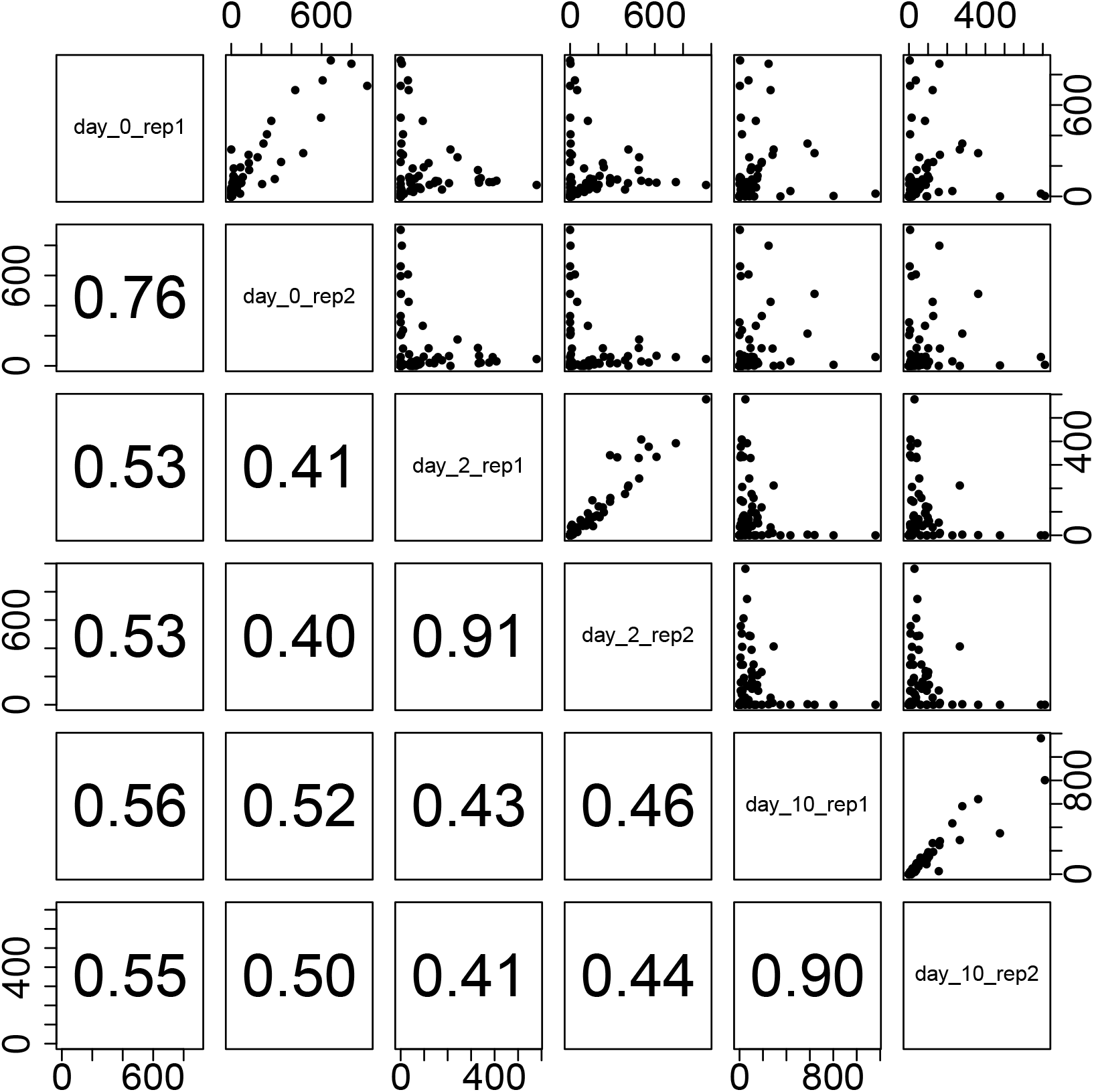
Upper: scatter plot of the number of single cells in Fig. 9, *N_IJ_*, 1 ≤ *I,J* ≥ 10, in one of 10 × 10 regions, *S_IJ_*. Lower: Kendall’s’s correlation coefficients of *N_IJ_* between six samples.

**Figure 11:**
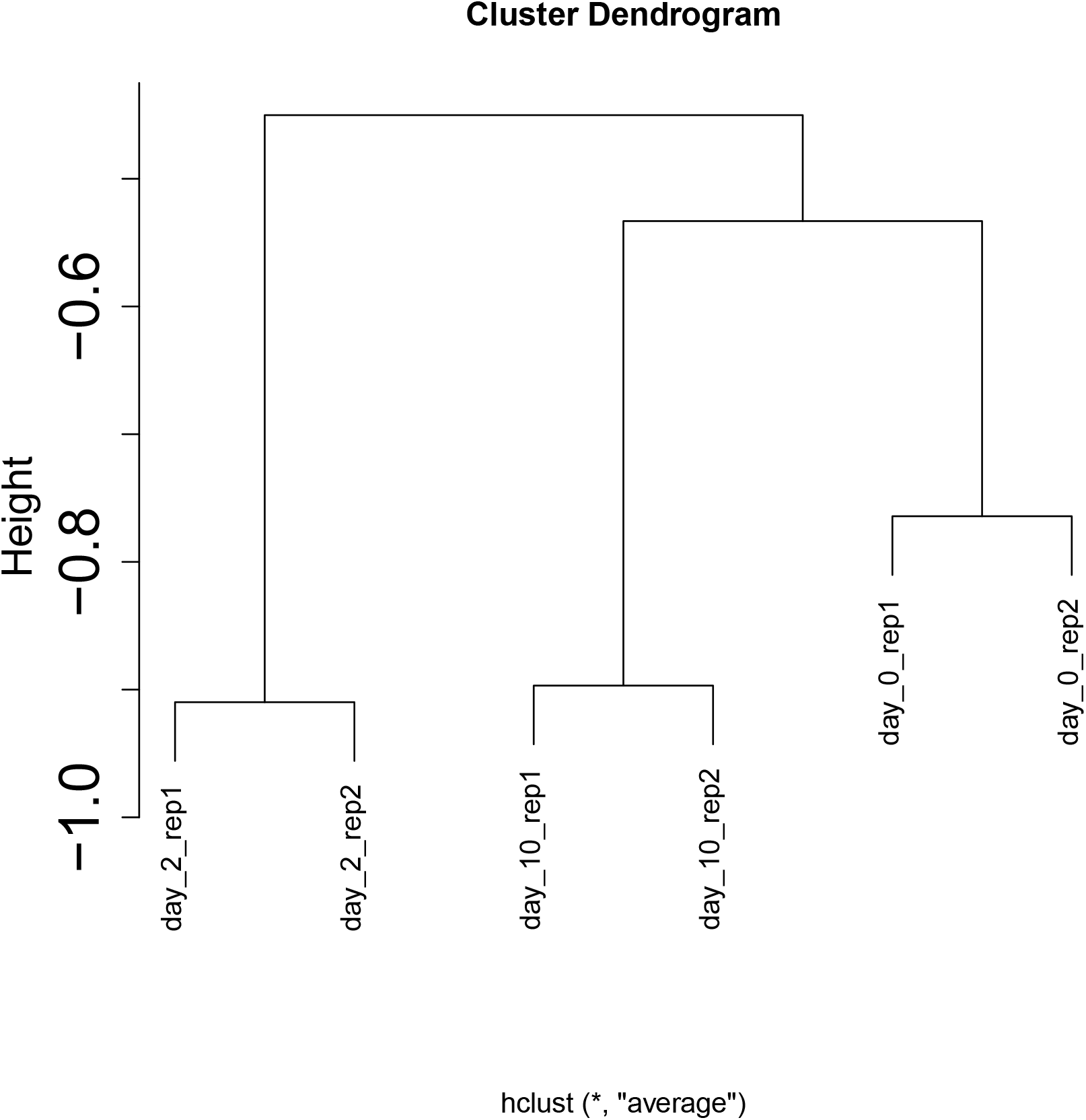
Hierarchical clustering by UPGMA of six samples (Table 2) using coordinates obtained by UMAP. Distance is represented by negatively signed correlation coefficients in Fig. 10

## 5. Conclusions

In this paper, we applied TD to an scATAC-seq dataset, and the obtained embedding can be used for UMAP, after which the embedded material obtained by UMAP can differentiate tissues from which the scATAC sequence was retrieved. TD can work with large sparse datasets generated by approximately 200 bp intervals, as these can be stored in a sparse matrix format. The large size of these datasets cannot be processed by any other method. The proposed method is the only method that can work with high-resolution native scATAC-seq datasets.

## Funding

This work was supported by KAKENHI (Grant Numbers 20K12067) to Y.-H.T.

## CRediT authorship contribution statement

**Y-h. Taguchi:** Y.-H.T. planned the research and performed the analyses. Y.-H.T. evaluated the results, discussions, and outcomes and wrote and reviewed the manuscript and has read and agreed to the published version of the manuscript.. **Turki Turki:** T.T. evaluated the results, discussions, and outcomes and wrote and reviewed the manuscript and has read and agreed to the published version of the manuscript..

---

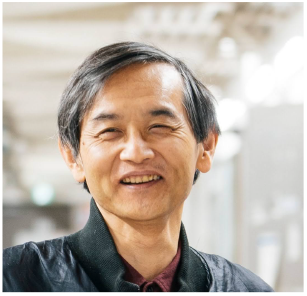
Y-H. TAGUCHI received a BS in Physics from the Tokyo Institute of Technology and a PhD in Physics from the Tokyo Institute of Technology. He is currently a full professor with the Department of Physics, Chuo University, Japan. His works have been published in leading journals such as Physical Review Letters, Bioinformatics, and Scientific Reports. His research interests include bioinformatics, machine learning, and nonlinear physics. He is also an editorial board member of PloS ONE, BMC Medical Genomics, Medicine (Lippincott Williams & Wilkins journal), BMC Research Notes, non-coding RNA (MDPI), and IPSJ Transaction on Bioinformatics.

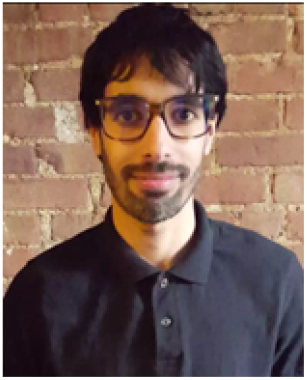
TURKITURKI has a BS in Computer Science from King Abdulaziz University, an MS in Computer Science from New York University Tandon School of Engineering, and a PhD in Computer Science from the New Jersey Institute of Technology. He is currently an associate professor with the Department of Computer Science, King Abdulaziz University, Saudi Arabia. His research interests include machine learning, bioinformatics, and computational genomics. His work has been published in journals such as Scientific Reports, Computers in Biology and Medicine, and Knowledge-Based Systems. He is also an editorial board member of PLoS ONE, BMC Medical Genomics, Computers in Biology and Medicine, and Sustainable Computing: Informatics and Systems.

## Notes

https://github.com/tagtag/scATAC-seq

### Competing Interest Statement

The authors have declared no competing interest.

### Summary of Updates

Title Changed

## References

[1] Baek, S., Lee, I., 2020. Single-cell ATAC sequencing analysis: From data preprocessing to hypothesis generation. Computational and Structural Biotechnology Journal 18, 1429–1439. URL:https://www.sciencedirect.com/science/article/pii/S2001037020303019, doi:https://doi.org/10.1016/j.csbj.2020.06.012.

[2] Baglama, J., Reichel, L., Lewis, B.W., 2021. irlba: Fast Truncated Singular Value Decomposition and Principal Components Analysis for Large Dense and Sparse Matrices. URL:https://CRAN.R-project.org/package=irlba. r package version 2.3.5.

[3] Barrett, T., Wilhite, S.E., Ledoux, P., Evangelista, C., Kim, I.F., Tomashevsky, M., Marshall, K.A., Phillippy, K.H., Sherman, P.M., Holko, M., Yefanov, A., Lee, H., Zhang, N., Robertson, C.L., Serova, N., Davis, S., Soboleva, A., 2012. NCBI GEO: archive for functional genomics data sets-update. Nucleic Acids Research 41, D991–D995. URL:https://doi.org/10.1093/nar/gks1193, doi:10.1093/nar/gks1193, arXiv:https://academic.oup.com/nar/article-pdf/41/D1/D991/3678141/gks1193.pdf.

[4] Buenrostro, J.D., Wu, B., Litzenburger, U.M., Ruff, D., Gonzales, M.L., Snyder, M.P., Chang, H.Y., Greenleaf, W.J., 2015. Single-cell chromatin accessibility reveals principles of regulatory variation. Nature 523, 486–490.

[5] Cavalcante, R.G., Sartor, M.A., 2017. annotatr: genomic regions in context. Bioinformatics 33, 2381–2383. URL: https://doi.org/10.1093/bioinformatics/btx183, doi:10.1093/bioinformatics/btx183, arXiv:https://academic.oup.com/bioinformatics/article-pdf/33/15/2381/25157896/btx183.pdf.

[6] Chen, H., Lareau, C., Andreani, T., Vinyard, M.E., Garcia, S.P., Clement, K., Andrade-Navarro, M.A., Buenrostro, J.D., Pinello, L., 2019. Assessment of computational methods for the analysis of single-cell ATAC-seq data. Genome Biology 20. URL:https://doi.org/10.1186/s13059-019-1854-5, doi:10.1186/s13059-019-1854-5.

[7] Fang, R., Preissl, S., Li, Y., Hou, X., Lucero, J., Wang, X., Motamedi, A., Shiau, A.K., Zhou, X., Xie, F., Mukamel, E.A., Zhang, K., Zhang, Y., Behrens, M.M., Ecker, J.R., Ren, B., 2021. Comprehensive analysis of single cell ATAC-seq data with SnapATAC. Nature Communications 12. URL:https://doi.org/10.1038/s41467-021-21583-9, doi:10.1038/s41467-021-21583-9.

[8] Giansanti, V., Tang, M., Cittaro, D., 2020. Fast analysis of scatac-seq data using a predefined set of genomic regions [version 2; peer review: 2 approved]. F1000Research 9. doi:10.12688/f1000research.22731.2.

[9] Grandi, F.C., Modi, H., Kampman, L., Corces, M.R., 2022. Chromatin accessibility profiling by ATAC-seq. Nature Protocols 17, 1518–1552. URL:https://doi.org/10.1038/s41596-022-00692-9, doi:10.1038/s41596-022-00692-9.

[10] Granja, J.M., Corces, M.R., Pierce, S.E., Bagdatli, S.T., Choudhry, H., Chang, H.Y., Greenleaf, W.J., 2021. ArchR is a scalable software package for integrative single-cell chromatin accessibility analysis. Nat Genet 53, 403–411.

[11] Guo, H., Yang, Z., Jiang, T., Liu, S., Wang, Y., Cui, Z., 2022. Evaluation of classification in single cell atac-seq data with machine learning methods. BMC Bioinformatics 23. URL:https://doi.org/10.1186/s12859-022-04774-z, doi:10.1186/s12859-022-04774-z.

[12] Huber, W., Carey, V.J., Gentleman, R., Anders, S., Carlson, M., Carvalho, B.S., Bravo, H.C., Davis, S., Gatto, L., Girke, T., Gottardo, R., Hahne, F., Hansen, K.D., Irizarry, R.A., Lawrence, M., Love, M.I., MacDonald, J., Obenchain, V., Oleś, A.K., Pagès, H., Reyes, A., Shannon, P., Smyth, G.K., Tenenbaum, D., Waldron, L., Morgan, M., 2015. Orchestrating high-throughput genomic analysis with bioconductor. Nature Methods 12, 115–121. URL:https://doi.org/10.1038/nmeth.3252, doi:10.1038/nmeth.3252.

[13] Konopka, T., 2022. umap: Uniform Manifold Approximation and Projection. URL:https://CRAN.R-project.org/package=umap. r package version 0.2.8.0.

[14] Kopp, W., Akalin, A., Ohler, U., 2022. Simultaneous dimensionality reduction and integration for single-cell ATAC-seq data using deep learning. Nat Mach Intell 4, 162–168.

[15] Kuleshov, M.V., Jones, M.R., Rouillard, A.D., Fernandez, N.F., Duan, Q., Wang, Z., Koplev, S., Jenkins, S.L., Jagodnik, K.M., Lachmann, A., McDermott, M.G., Monteiro, C.D., Gundersen, G.W., Ma’ayan, A., 2016. Enrichr: a comprehensive gene set enrichment analysis web server 2016 update. Nucleic Acids Research 44, W90–W97. URL:https://doi.org/10.1093/nar/gkw377, doi:10.1093/nar/gkw377, arXiv:https://academic.oup.com/nar/article-pdf/44/W1/W90/18788036/gkw377.pdf.

[16] Lee, D.R., Rhodes, C., Mitra, A., Zhang, Y., Maric, D., Dale, R.K., Petros, T.J., 2022. Transcriptional heterogeneity of ventricular zone cells in the ganglionic eminences of the mouse forebrain. eLife 11, e71864. URL:https://doi.org/10.7554/eLife.71864, doi: 10.7554/eLife.71864.

[17] Li, Z., Kuppe, C., Ziegler, S., Cheng, M., Kabgani, N., Menzel, S., Zenke, M., Kramann, R., Costa, I.G., 2021a. Chromatin-accessibility estimation from single-cell ATAC-seq data with scOpen. Nature Communications 12. URL:https://doi.org/10.1038/s41467-021-26530-2, doi:10.1038/s41467-021-26530-2.

[18] Li, Z., Kuppe, C., Ziegler, S., Cheng, M., Kabgani, N., Menzel, S., Zenke, M., Kramann, R., Costa, I.G., 2021b. Chromatin-accessibility estimation from single-cell ATAC-seq data with scOpen. Nat Commun 12, 6386.

[19] Mitchel, J., Gordon, M.G., Perez, R.K., Biederstedt, E., Bueno, R., Ye, C.J., Kharchenko, P.V., 2022. Tensor decomposition reveals coordinated multicellular patterns of transcriptional variation that distinguish and stratify disease individuals. bioRxiv URL: https://www.biorxiv.org/content/early/2022/02/18/2022.02.16.480703, doi:10.1101/2022.02.16.480703, arXiv:https://www.biorxiv.org/content/early/2022/02/18/2022.02.16.480703.full.pdf.

[20] Pan, X., Li, Z., Qin, S., Yu, M., Hu, H., 2021. ScLRTC: imputation for single-cell RNA-seq data via low-rank tensor completion. BMC Genomics 22. URL:https://doi.org/10.1186/s12864-021-08101-3, doi:10.1186/s12864-021-08101-3.

[21] R Core Team, 2022. R: A Language and Environment for Statistical Computing. R Foundation for Statistical Computing. Vienna, Austria. URL:https://www.R-project.org/.

[22] Satpathy, A.T., Granja, J.M., Yost, K.E., Qi, Y., Meschi, F., McDermott, G.P., Olsen, B.N., Mumbach, M.R., Pierce, S.E., Corces, M.R., Shah, P., Bell, J.C., Jhutty, D., Nemec, C.M., Wang, J., Wang, L., Yin, Y., Giresi, P.G., Chang, A.L.S., Zheng, G.X.Y., Greenleaf, W.J., Chang, H.Y., 2019. Massively parallel single-cell chromatin landscapes of human immune cell development and intratumoral T cell exhaustion. Nat Biotechnol 37, 925–936.

[23] Song, Q., Zhu, X., Jin, L., Chen, M., Zhang, W., Su, J., 2022. SMGR: a joint statistical method for integrative analysis of single-cell multi-omics data. NAR Genomics and Bioinformatics 4. URL:https://doi.org/10.1093/nargab/lqac056, doi:10.1093/nargab/lqac056.

[24] Stuart, T., Butler, A., Hoffman, P., Hafemeister, C., Papalexi, E., Mauck, W.M., Hao, Y., Stoeckius, M., Smibert, P., Satija, R., 2019. Comprehensive integration of single-cell data. Cell 177, 1888–1902.e21. URL:https://www.sciencedirect.com/science/article/pii/S0092867419305598, doi:https://doi.org/10.1016/j.cell.2019.05.031.

[25] Stuart, T., Srivastava, A., Madad, S., Lareau, C., Satija, R., 2021. Single-cell chromatin state analysis with signac. Nature Methods URL: https://doi.org/10.1038/s41592-021-01282-5, doi:10.1038/s41592-021-01282-5.

[26] Taguchi, Y.H., 2020. Unsupervised Feature Extraction Applied to Bioinformatics. Springer International Publishing. URL:https://doi.org/10.1007/978-3-030-22456-1, doi:10.1007/978-3-030-22456-1.

[27] Taguchi, Y.h., Turki, T., 2021. Tensor-decomposition-based unsupervised feature extraction in single-cell multiomics data analysis. Genes 12. URL:https://www.mdpi.com/2073-4425/12/9/1442, doi:10.3390/genes12091442.

[28] Xiong, L., Xu, K., Tian, K., Shao, Y., Tang, L., Gao, G., Zhang, M., Jiang, T., Zhang, Q.C., 2019. SCALE method for single-cell ATAC-seq analysis via latent feature extraction. Nat Commun 10, 4576.

[29] Xu, Y., Begoli, E., McCord, R.P., 2022. sciCAN: single-cell chromatin accessibility and gene expression data integration via cycle-consistent adversarial network. npj Systems Biology and Applications 8. URL:https://doi.org/10.1038/s41540-022-00245-6, doi:10.1038/s41540-022-00245-6.

[30] Yuan, H., Kelley, D.R., 2022. scBasset: sequence-based modeling of single-cell ATAC-seq using convolutional neural networks. Nature Methods 19, 1088–1096. URL:https://doi.org/10.1038/s41592-022-01562-8, doi:10.1038/s41592-022-01562-8.

