## Supplementary material for "Tensor Decomposition Discriminates Tissues Using scATAC-seq": Supp_Tables.pdf

### Supplementary tables

Table S1: Details of number of annotations in pie chart shown in Fig. ??

| | |
|---|---|
| genes_intergenic | genes_3UTRs |
| 909 | 1097 |
| enhancers_fantom | cpg_shelves |
| 1388 | 1772 |
| cpg_inter | genes_cds |
| 2250 | 4925 |
| cpg_islands | genes_5UTRs |
| 5555 | 5565 |
| lncrna_gencode | cpg_shores |
| 8806 | 9054 |
| genes_firstexons | genes_intronexonboundaries |
| 10218 | 13052 |
| genes_exons | genes_exonintronboundaries |
| 15803 | 16799 |
| genes_promoters | genes_1to5kb |
| 22589 | 29173 |
| genes_introns | |
| 39852 | |

Table S2: Top 10 TF in "ENCODE and ChEA Consensus TFs from ChIP-X" of Enrichr

| Term | Overlap | P-value | Adjusted P-value |
|---|---|---|---|
| PBX3 ENCODE | 238/1269 | $3.42 \times 10^{-64}$ | $3.55 \times 10^{-62}$ |
| IRF3 ENCODE | 158/663 | $3.81 \times 10^{-56}$ | $1.98 \times 10^{-54}$ |
| NFYB ENCODE | 429/3715 | $3.78 \times 10^{-54}$ | $1.31 \times 10^{-52}$ |
| NFYA ENCODE | 312/2250 | $7.23 \times 10^{-54}$ | $1.88 \times 10^{-52}$ |
| FOS ENCODE | 148/637 | $5.10 \times 10^{-51}$ | $1.06 \times 10^{-49}$ |
| SP1 ENCODE | 153/707 | $1.53 \times 10^{-48}$ | $2.65 \times 10^{-47}$ |
| SP2 ENCODE | 168/994 | $2.36 \times 10^{-38}$ | $3.51 \times 10^{-37}$ |
| CREB1 CHEA | 205/1444 | $1.74 \times 10^{-35}$ | $2.26 \times 10^{-34}$ |
| RFX5 ENCODE | 104/559 | $3.46 \times 10^{-27}$ | $4.00 \times 10^{-26}$ |
| UBTF ENCODE | 183/1631 | $2.20 \times 10^{-19}$ | $2.28 \times 10^{-18}$ |

Table S3: Top 10 TF in "ENCODE TF ChIP-seq 2015" of Enrichr

| Term | Overlap | P-value | Adjusted P-value |
|---|---|---|---|
| IRF3 HepG2 hg19 | 162/755 | $6.11 \times 10^{-51}$ | $4.98 \times 10^{-48}$ |
| IRF3 HeLa-S3 hg19 | 264/1809 | $6.24 \times 10^{-49}$ | $2.55 \times 10^{-46}$ |
| FOS GM12878 hg19 | 210/1244 | $2.51 \times 10^{-48}$ | $6.83 \times 10^{-46}$ |
| SP1 K562 hg19 | 203/1249 | $5.26 \times 10^{-44}$ | $1.07 \times 10^{-41}$ |
| CHD2 MEL cell line mm9 | 255/1826 | $1.52 \times 10^{-43}$ | $2.10 \times 10^{-41}$ |
| FOS K562 hg19 | 270/2000 | $1.54 \times 10^{-43}$ | $2.10 \times 10^{-41}$ |
| SP2 H1-hESC hg19 | 185/1184 | $1.64 \times 10^{-37}$ | $1.91 \times 10^{-35}$ |
| SP1 HCT116 hg19 | 219/1577 | $1.45 \times 10^{-36}$ | $1.47 \times 10^{-34}$ |
| SP2 HepG2 hg19 | 211/1507 | $1.11 \times 10^{-35}$ | $1.01 \times 10^{-33}$ |
| FOS HeLa-S3 hg19 | 146/860 | $1.85 \times 10^{-33}$ | $1.51 \times 10^{-31}$ |

Table S4: Top 10 TF in "TRANSFAC and JASPAR PWMs" of Enrichr

| Term | Overlap | P-value | Adjusted P-value |
|---|---|---|---|
| SP1 (mouse) | 242/2360 | $1.56 \times 10^{-20}$ | $4.84 \times 10^{-18}$ |
| PCBP1 (human) | 142/1360 | $1.26 \times 10^{-12}$ | $1.96 \times 10^{-10}$ |
| SP1 (human) | 141/1406 | $2.95 \times 10^{-11}$ | $3.05 \times 10^{-9}$ |
| TEAD2 (mouse) | 154/1591 | $4.74 \times 10^{-11}$ | $3.67 \times 10^{-9}$ |
| TEAD4 (human) | 127/1354 | $1.93 \times 10^{-8}$ | $1.20 \times 10^{-6}$ |
| TCFAP2A (human) | 126/1367 | $6.01 \times 10^{-8}$ | $2.92 \times 10^{-6}$ |
| SMAD4 (mouse) | 141/1580 | $6.60 \times 10^{-8}$ | $2.92 \times 10^{-6}$ |
| SP3 (human) | 121/1332 | $2.47 \times 10^{-7}$ | $8.98 \times 10^{-6}$ |
| EGR1 (mouse) | 141/1617 | $2.61 \times 10^{-7}$ | $8.98 \times 10^{-6}$ |
| E2F6 (human) | 114/1358 | $2.26 \times 10^{-5}$ | $6.99 \times 10^{-4}$ |

Table S5: Top 10 experiments in "Allen Brain Atlas 10x scRNA 2021" of Enrichr

| Term | Overlap | P-value | Adjusted P-value |
|---|---|---|---|
| Mouse 359 OPC down | 96/841 | $6.78 \times 10^{-11}$ | $4.30 \times 10^{-8}$ |
| Human Endo L2-5 NOSTRIN SRGN down | 83/703 | $2.76 \times 10^{-10}$ | $8.75 \times 10^{-8}$ |
| Mouse 372 SMC down | 62/466 | $5.24 \times 10^{-10}$ | $8.77 \times 10^{-8}$ |
| Mouse 375 VLMC down | 68/535 | $5.52 \times 10^{-10}$ | $8.77 \times 10^{-8}$ |
| Mouse 357 Astro down | 77/651 | $1.15 \times 10^{-9}$ | $1.46 \times 10^{-7}$ |
| Human Astro L1-6 FGFR3 PLCG1 down | 95/880 | $1.67 \times 10^{-9}$ | $1.77 \times 10^{-7}$ |
| Mouse 356 Astro down | 73/617 | $3.10 \times 10^{-9}$ | $2.67 \times 10^{-7}$ |
| Human VLMC L1-5 PDGFRA COLEC12 down | 83/742 | $3.68 \times 10^{-9}$ | $2.67 \times 10^{-7}$ |
| Mouse 374 VLMC down | 64/513 | $3.78 \times 10^{-9}$ | $2.67 \times 10^{-7}$ |
| Mouse 377 Micro down | 50/365 | $9.54 \times 10^{-9}$ | $6.06 \times 10^{-7}$ |

Table S6: Top 10 cells in "CellMarker Augmented 2021" of Enrichr

| Term | Overlap | P-value | Adjusted P-value |
|---|---|---|---|
| Radial Glial cell:Undefined | 6/11 | $1.26 \times 10^{-5}$ | $8.04 \times 10^{-3}$ |
| Neural Stem cell:Brain | 8/25 | $5.14 \times 10^{-5}$ | $1.46 \times 10^{-2}$ |
| Neural Stem cell:Undefined | 12/58 | $9.15 \times 10^{-5}$ | $1.46 \times 10^{-2}$ |
| Purkinje cell:Brain | 16/96 | $1.06 \times 10^{-4}$ | $1.46 \times 10^{-2}$ |
| Natural Killer T (NKT) cell:Fetal Kidney | 312/4543 | $1.41 \times 10^{-4}$ | $1.46 \times 10^{-2}$ |
| Mesoderm cell:Undefined | 16/99 | $1.55 \times 10^{-4}$ | $1.46 \times 10^{-2}$ |
| Astrocyte:Embryonic Prefrontal Cortex | 37/342 | $1.60 \times 10^{-4}$ | $1.46 \times 10^{-2}$ |
| Cancer Stem cell:Brain | 8/30 | $2.15 \times 10^{-4}$ | $1.71 \times 10^{-2}$ |
| Pancreatic Polypeptide cell:Pancreas | 6/18 | $3.59 \times 10^{-4}$ | $2.53 \times 10^{-2}$ |
| Neural Progenitor cell:Embryonic Prefrontal Cortex | 21/166 | $5.48 \times 10^{-4}$ | $2.74 \times 10^{-2}$ |

Table S7: Top 10 terms in "GO Biological Process 2021" of Enrichr

| Term | Overlap | P-value | Adjusted P-value |
|---|---|---|---|
| negative regulation of transcription, DNA-templated (GO:0045892) | 117/948 | $1.90 \times 10^{-15}$ | $6.51 \times 10^{-12}$ |
| positive regulation of transcription, DNA-templated (GO:0045893) | 123/1183 | $6.35 \times 10^{-11}$ | $1.09 \times 10^{-7}$ |
| negative regulation of transcription by RNA polymerase II (GO:0000122) | 82/684 | $1.68 \times 10^{-10}$ | $1.92 \times 10^{-7}$ |
| negative regulation of cellular macromolecule biosynthetic process (GO:2000113) | 69/547 | $5.72 \times 10^{-10}$ | $4.90 \times 10^{-7}$ |
| positive regulation of transcription by RNA polymerase II (GO:0045944) | 97/908 | $1.93 \times 10^{-9}$ | $1.33 \times 10^{-6}$ |
| negative regulation of nucleic acid-templated transcription (GO:1903507) | 59/464 | $7.67 \times 10^{-9}$ | $4.38 \times 10^{-6}$ |
| regulation of transcription by RNA polymerase II (GO:0006357) | 188/2206 | $1.04 \times 10^{-8}$ | $5.08 \times 10^{-6}$ |
| positive regulation of nucleic acid-templated transcription (GO:1903508) | 62/511 | $1.92 \times 10^{-8}$ | $8.22 \times 10^{-6}$ |
| regulation of transcription, DNA-templated (GO:0006355) | 184/2244 | $2.44 \times 10^{-7}$ | $9.30 \times 10^{-5}$ |
| negative regulation of neuron differentiation (GO:0045665) | 10/24 | $3.47 \times 10^{-7}$ | $1.19 \times 10^{-4}$ |

Table S8: Top 10 terms in "GO Molecular Function 2021" of Enrichr

| Term | Overlap | P-value | Adjusted P-value |
|---|---|---|---|
| sequence-specific double-stranded DNA binding (GO:1990837) | 81/712 | $2.63 \times 10^{-9}$ | $1.61 \times 10^{-6}$ |
| sequence-specific DNA binding (GO:0043565) | 78/707 | $2.02 \times 10^{-8}$ | $6.17 \times 10^{-6}$ |
| double-stranded DNA binding (GO:0003690) | 71/651 | $1.39 \times 10^{-7}$ | $2.83 \times 10^{-5}$ |
| DNA-binding transcription factor binding (GO:0140297) | 29/208 | $8.46 \times 10^{-6}$ | $1.29 \times 10^{-3}$ |
| transcription regulatory region nucleic acid binding (GO:0001067) | 29/212 | $1.23 \times 10^{-5}$ | $1.50 \times 10^{-3}$ |
| DNA binding (GO:0003677) | 74/811 | $5.08 \times 10^{-5}$ | $5.18 \times 10^{-3}$ |
| acetylation-dependent protein binding (GO:0140033) | 7/21 | $1.14 \times 10^{-4}$ | $8.75 \times 10^{-3}$ |
| lysine-acetylated histone binding (GO:0070577) | 7/21 | $.14 \times 10^{-4}$ | $8.75 \times 10^{-3}$ |
| dihydropyrimidinase activity (GO:0004157) | 4/6 | $1.47 \times 10^{-4}$ | $1.00 \times 10^{-2}$ |
| mRNA binding (GO:0003729) | 30/263 | $2.62 \times 10^{-4}$ | $1.61 \times 10^{-2}$ |
